# Pan-cancer molecular signatures connecting aspartate transaminase (AST) to cancer prognosis, metabolic and immune signatures

**DOI:** 10.1101/2024.03.01.582939

**Authors:** Geoffrey H. Siwo, Amit G. Singal, Akbar K. Waljee

## Abstract

**Background:** Serum aspartate transaminase (sAST) level is used routinely in conjunction with other clinical assays to assess liver health and disease. Increasing evidence suggests that sAST is associated with all-cause mortality and has prognostic value in several cancers, including gastrointestinal and urothelial cancers. Here, we undertake a systems approach to unravel molecular connections between AST and cancer prognosis, metabolism, and immune signatures at the transcriptomic and proteomic levels.

**Methods:** We mined public gene expression data across multiple normal and cancerous tissues using the Genotype Tissue Expression (GTEX) resource and The Cancer Genome Atlas (TCGA) to assess the expression of genes encoding AST isoenzymes (GOT1 and GOT2) and their association with disease prognosis and immune infiltration signatures across multiple tumors. We examined the associations between AST and previously reported pan-cancer molecular subtypes characterized by distinct metabolic and immune signatures. We analyzed human protein-protein interaction networks for interactions between GOT1 and GOT2 with cancer-associated proteins. Using public databases and protein-protein interaction networks, we determined whether the subset of proteins that interact with AST (GOT1 and GOT2 interactomes) are enriched with proteins associated with specific diseases, miRNAs and transcription factors.

**Results:** We show that AST transcript isoforms (GOT1 and GOT2) are expressed across a wide range of normal tissues. AST isoforms are upregulated in tumors of the breast, lung, uterus, and thymus relative to normal tissues but downregulated in tumors of the liver, colon, brain, kidney and skeletal sarcomas. At the proteomic level, we find that the expression of AST is associated with distinct pan-cancer molecular subtypes with an enrichment of specific metabolic and immune signatures. Based on human protein-protein interaction data, AST physically interacts with multiple proteins involved in tumor initiation, suppression, progression, and treatment. We find enrichments in the AST interactomes for proteins associated with liver and lung cancer and dermatologic diseases. At the regulatory level, the GOT1 interactome is enriched with the targets of cancer-associated miRNAs, specifically mir34a – a promising cancer therapeutic, while the GOT2 interactome is enriched with proteins that interact with cancer-associated transcription factors.

**Conclusions:** Our findings suggest that perturbations in the levels of AST within specific tissues reflect pathophysiological changes beyond tissue damage and have implications for cancer metabolism, immune infiltration, prognosis, and treatment personalization.

## Introduction

The measurement of circulating aspartate transaminase enzyme levels in serum (sAST) is commonly performed in clinical laboratories to assess liver health and damage. In humans, AST occurs as two isoenzymes encoded by distinct genes: a cytoplasmic (Glutamic Oxaloacetic Transaminase 1, GOT1) and a mitochondrial isoform (GOT2) ^6^. GOT1 is expressed mainly in red blood cells and the heart, while GOT2 is expressed predominantly in the liver^6^. Clinical laboratory tests for AST measure its level in serum (sAST) and do not differentiate between the two isoenzymes^6^.

Even though AST is typically used in the context of liver damage, the enzyme is expressed in a wide range of tissues, including in the gastrointestinal system, skeletal muscle, heart, and kidneys ^7^. High levels of sAST have been associated with various liver diseases including cirrhosis, metabolic dysfunction associated steatotic liver disease (MASLD) and hepatocellular carcinoma (HCC) ^8–10^. In addition, sAST has been associated with several non-liver tumors, cardiovascular diseases, type 2 diabetes and all-cause mortality^2,3,11^. Besides these epidemiological associations, increasing evidence from molecular studies paint a complex role of AST isoenzymes in cancer ^12–16^, encephalopathies ^17,18^, inborn errors of metabolism^19^, ischemic stroke ^20^ and neurodegenerative diseases such as Parkinson’s, Alzheimer’s, and Huntington’s disease ^21^. Epidemiologically, sAST has been associated with prognosis of cancers spanning multiple tissues including non-metastatic renal cell carcinoma ^22,23^, head and neck cancer ^24^, gastric adenocarcinoma ^25^, esophageal squamous cell adenocarcinoma ^26^, upper urinary tract urothelial carcinoma ^27^, prostate cancer ^28^, and Pancreatic Ductal Adenocarcinoma (PDAC) ^29^. While these studies have provided evidence for prognostic associations between sAST and cancer, insights into the underlying biological mechanisms are lacking.

Metabolic reprogramming, the process by which cancer cells rewire their metabolism to support the high demand for proteins, nucleotides and lipids required for rapid growth and proliferation is increasingly considered a hallmark of cancer ^30^ The non-essential amino acid glutamine ^31^ is critical for cancer cell growth and proliferation ^32,33^. Glutamine is broken down into glutamate by the action of glutaminases and the resulting glutamate can be used to generate α-ketoglutarate. AST catalyzes the reversible conversion of glutamate to aspartate which provides a carbon and nitrogen source for the biosynthesis of purines, pyrimidines, amino acids and driving the malate-aspartate shuttle for redox homeostasis ^34^. Consequently, in some tumors, growth and proliferation are highly dependent on aspartate ^35–37^. GOT1 catalyzes the conversion of α-ketoglutarate and aspartate into oxaloacetate and glutamate while the GOT2 catalyzes the reverse reaction ^38^. In PDAC, GOT1 has been reported as a prognostic marker ^39^ and functionally linked to KRAS-dependent metabolic reprogramming ^40^. Further, inhibition of GOT1 was found to impair pancreatic cancer growth ^41–43^. GOT2 has been associated with cancer through several mechanisms^44^. For example, GOT2 can traffic fatty acids leading to suppression of antitumor immunity via the peroxisome proliferator activated receptor alpha-dependent pathway (PPAR) ^45,46^. In triple negative breast cancer, GOT2 overexpression was associated with increased proliferation ^47–49^, while in clear cell renal cell carcinoma GOT2 was downregulated and associated with poor survival and immune infiltrates ^50^. A gain of function mutation in GOT2 has been reported in rare neurological tumors that also harbor loss of function mutations in the malate aspartate shuttle ^51,52^. Exogenous expression of GOT2 in chimeric antigen receptor (CAR)-T cells enhanced in vitro and in vivo cytotoxic activity in liver cancer ^53^. Collectively, these studies indicate that the associations between AST and various cancers at the epidemiological level may be driven by several molecular mechanisms.

To date, there have been no comprehensive studies involving pan-tissue transcriptomic and proteomic assessments of AST isoforms across several cancers and their association with tumor prognosis or metabolic and immune infiltration. In this study, we leverage public genomic and proteomic datasets to investigate pan-cancer associations between GOT1 and GOT2 gene expression and protein levels with tumor prognosis, metabolic and immune signatures.

## Methods

### Pan-tissue, Pan-cancer transcriptome analyses of AST levels

The expression of genes encoding AST isoenzymes (GOT1 and GOT2) across tissues was performed using data from the Genotype Tissue Expression portal (GTEX Analysis Release V8) based on samples from 54 non-diseased tissues of nearly 1000 individuals ^54^. Data was visualized using boxplots on the GTEX portal with transcript levels of GOT1 and GOT2 represented as Transcripts per Million (TPM). Only bulk gene expression was considered. The expression level of each was based on a gene model in which, alternatively, spliced transcript isoforms are collapsed into single genes. The transcript levels of GOT1 and GOT2 in cancer cells were based on tumor and the corresponding normal tissue data from The Cancer Genome Atlas (TCGA) ^55^. Analyses and visualization of TCGA datasets were performed using the UALCAN resource ^56,57^. The comparisons and P-value estimates of expression level differences of GOT1 and GOT2 in normal vs tumor tissue were performed using UCSC Xena platform ^58^.

### Pan-cancer proteomic analyses of AST levels

We leveraged previously described cancer molecular subtypes based on proteogenomic characterization of 2002 primary tumors from 14 cancers types^59^ and proteomic datasets across 532 cancers spanning 6 tissue-based types (breast, colon, ovarian, renal and uterine)^60^. These subtypes were obtained previously from unsupervised clustering of the proteomic data of 2002 primary tumors leading to the identification of 11 molecular subtypes (s1 to s11), each including multiple tissue-based cancers ^59^. In addition, we leveraged ten molecular subtypes (k1 to k10) that were previously described and found to be enriched with distinct oncogenic and metabolic pathways ^60^. To determine whether the tumors expressing high or low protein levels of GOT1 and GOT2 are associated with distinct tumor molecular subtypes, we used the UALCAN portal ^56,57^ leveraging mass-spectrometry proteomic data across the previously described pan-cancer subtypes from the Clinical Proteomic Tumor Analysis Consortium (CPTAC) Confirmatory/ Discovery cohorts ^61^.

### Analyses of AST associations to cancer prognosis and immune infiltration signatures

Survival analysis was performed using TCGA data based on tumor gene expression levels of GOT1 and GOT2, separately from the Survival Genie portal ^62^. Tumors were categorized into those expressing high vs. low levels of GOT1 (GOT2) using the median gene expression value of the transcript derived from the RNA-seq normalized FPKM gene expression. The categories were then used to define tumor cohorts into high vs. low expressing groups for survival analyses using Kaplan Meir plots on Survival Genie ^62^. A P-value threshold of P <0.05 was used to define the significant association between GOT1 and GOT2 expression and overall survival. To determine correlations between GOT1 expression (or GOT2 separately) and tumor-infiltrating lymphocytes (TILs), an implementation of the CIBERSORT algorithm ^63,64^ within Survival Genie^62^ was used. Briefly, for each tumor sample, the relative fraction of TIL subtypes was estimated for validated LM6 and LM22 immune cell signatures using bulk tumor FPKM gene expression data based on the CIBERSORT deconvolution method ^64^. The tumor’s inferred cell composition was then correlated to the expression of GOT1 or GOT2. Significant positive or negative correlation was assessed using a cutoff of P < 0.05.

### Analysis of AST protein-protein interaction networks

GOT1 and GOT2 protein-protein interaction networks were obtained from the BioGrid database version 4.4 ^65^. The analysis of disease genes enriched in each network was assessed using the DisGeNET database version 7.0, which contains over 1 million disease-gene associations involving 21,671 genes and 30,170 diseases, traits, and human phenotypes involving 369,554 variant-disease associations ^66^. The statistical significance of disease enrichments was based on a P and q-value estimate of <0.05, and only the top 10 significantly enriched diseases were considered. The enrichment of miRNA target genes in the GOT1 and GOT2 networks was determined using Enrichr ^67–69^ and miRTarBase, a curated database of experimentally determined microRNA-target interactions ^70,71^. A significance threshold of both P and q-value < 0.05 was applied to the miRNA enrichments, where the q-value refers to P-values corrected for false discovery rates to control for multiple hypothesis testing.

### Analysis of Regulatory Interactions

Two types of regulatory interactions associated with GOT1 and GOT2 protein-protein networks were assessed. The first type of regulatory interactions involved transcription factor protein-protein interactions to identify GOT1 and GOT2 interacting proteins that also interact with specific transcription factors. The protein-protein interaction networks for transcription factors were based on the literature ^67–69^ while GOT1 and GOT2 networks were from BioGrid version 4.4 ^65^. For each transcription factor, the statistical significance of enrichment of proteins in its network was estimated using a P-value and corrected for multiple hypothesis testing using Benjamini-Hochberg method to generate a q-value. Enrichments were estimated using Enrichr ^67–69^ and only those with P- and q-values < 0.05 were regarded as significant.

## Results

### Pan-tissue, Pan-cancer expression of AST

To assess the expression level of AST across multiple tissues, we queried the GTEX resource. GOT1 and GOT2 expression was evident in multiple tissues with the highest levels of expression in skeletal muscle, the heart, liver and specific regions of the brain (Fig. 1). The lowest expression level was in whole blood. Tissues expressing high levels of GOT1 also tended to express high levels of GOT2.

**Fig. 1.**
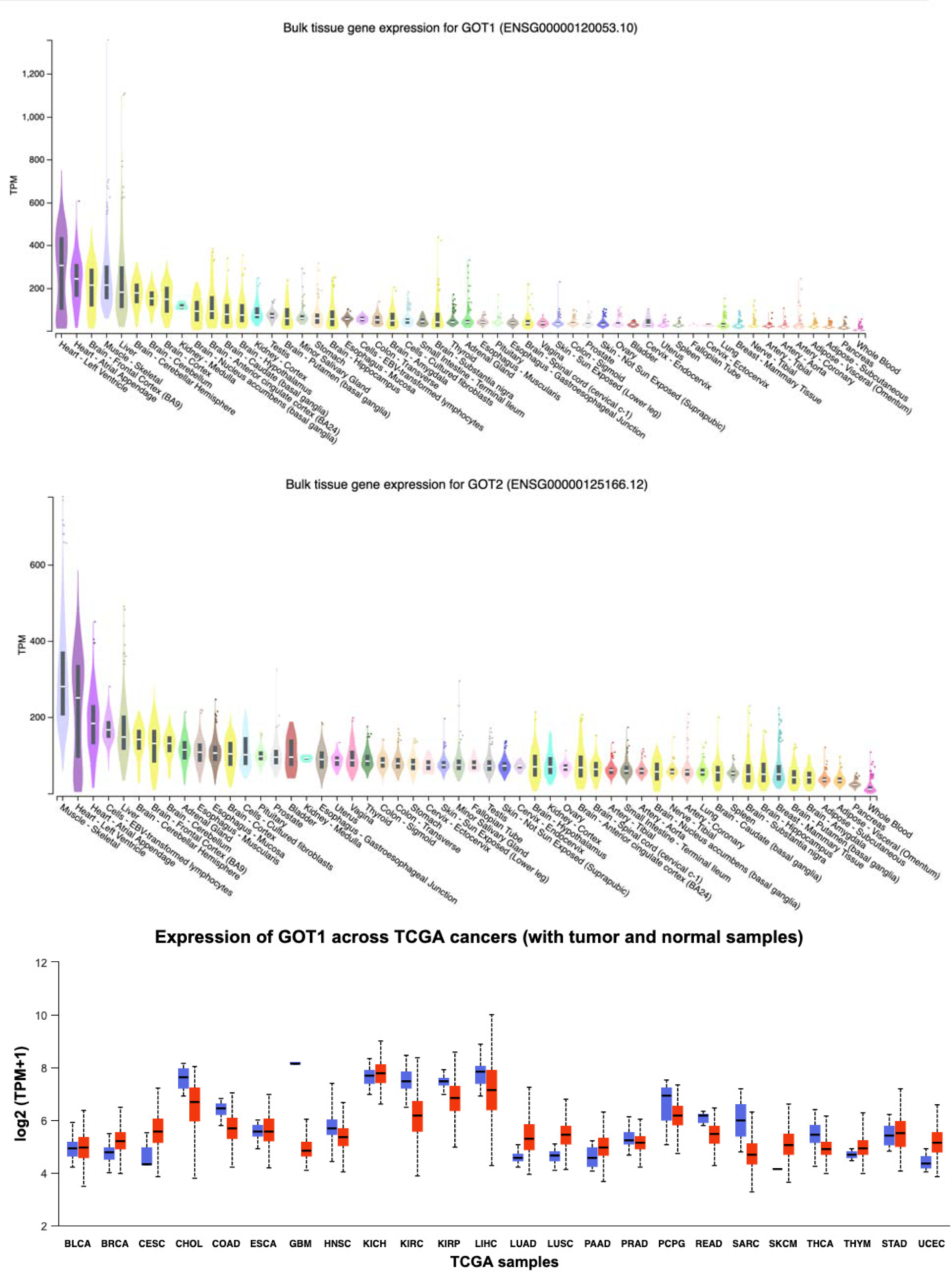
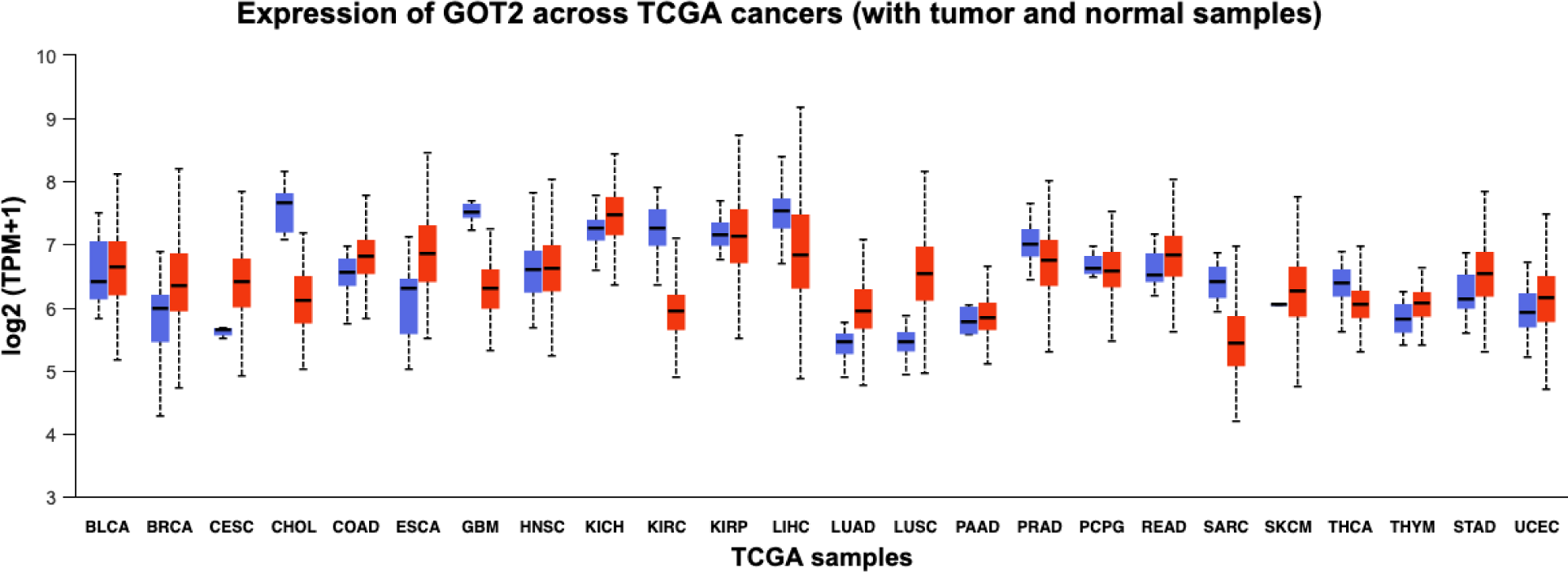
Pan-tissue and pan-cancer transcript levels of GOT1 and GOT2 based on GTEX and TCGA datasets, respectively.

Given the extensive pan-tissue expression of GOT1 and GOT2, we performed an analysis of their expression across multiple cancers in TCGA. Relative to the normal adjacent tissue, GOT1 expression was higher in breast cancer (BRCA, P << 0.001), cervical squamous cell carcinoma and endocervical adenocarcinoma (CESC, P = 0.12), lung adenocarcinoma (LUAD, P = 1.351e-13), Lung Squamous Cell Carcinoma (LUSC, P = 1.733e-24), Thymoma (THYM, P = 0.1559) and Uterine Corpus Endometrial Carcinoma (UCEC, P = 1.714e-12). However, GOT1 was expressed at lower levels in Cholangiocarcinoma (CHOL, P = 8.212e-11), Colorectal Adenocarcinoma (COAD, P = 1.998e-15), glioblastoma (GBM, P << 0.001), Head and Neck Squamous Cell Carcinoma (HNSC, P << 0.001), Kidney Renal Clear Cell Carcinoma (KIRC, P << 0.001), Kidney Renal Papillary Cell Carcinoma (KIRP, P = 0.0004776), Hepatocellular Carcinoma (LIHC, P = 4.127e-12), Pheochromocytoma and Paraganglioma (PCPG, P = 0.1990), Rectal Adenocarcinoma (READ, P = 4.797e-8), sarcoma (SARC, P = 0.003331) and Thyroid Carcinoma (THCA, P = 0.02) (Fig. 1).

Relative to the adjacent normal tissue, GOT2 expression was highly expressed in BRCA (P << 0.001), CESC (P = 0.1031), LUAD (P = 1.933e-13), LUSC (P = 4.948e-62), Pancreatic Adenocarcinoma (PAAD, P = 0.5721) and UCEC (P = 0.03423) but expressed at reduced levels in CHOL (P = 2.349e-17), GBM (P = 1.601e-9), KIRC (P << 0.001), LIHC (P = 1.110e-16), Prostate Adenocarcinoma (PRAD, P = 0.003360) and SARC (P = 0.005591). For both GOT1 and GOT2, tumors whose resident tissues normally have high expression of these genes tended to have low expression and vice versa. For example, while the liver and muscle are among the top tissues expressing high levels of GOT1 and GOT2 when compared to all other tissues in GTEX, tumors from these tissues had downregulated levels of expression of these genes relative to the corresponding normal tissue. Conversely, in lungs and uterus - tissues with low constitutive expression of GOT1 and GOT2-, these genes were upregulated in the tumor compared to the corresponding normal tissue.

### AST protein levels in pan-cancer proteomic subtypes

Given the extensive differential expression of GOT1 and GOT2 across multiple tumors coupled with prior studies linking the expression of these genes to tumor metabolic rewiring ^12,15,40,43,49,72,73^, we next investigated the association between GOT1 and GOT2 protein levels in a set of 10 (K1 to K10) and 11 (S1 to S11) previously described pan-cancer molecular subtypes across a set of 532 tumors ^60^ and additional set of 2002 tumors^59^ based on proteogenomic signatures.

The top pan-cancer subtypes associated with high expression of GOT1 were K5, K8 and K1, and S11 and S4 (Fig. 2, see Supplementary Table 1 for P-values). K5 subtypes were previously associated with tricarboxylic acid cycle (TCA) and oxidative phosphorylation as well as with targets of MYC, YAP1, and Ras pathway ^60^. K8 was associated with the Ras pathway, while K1 with the pentose phosphate pathway and glycolysis ^60^. S4 and S11 proteomic signatures were previously associated with increased activation of fatty acid metabolism, glycolysis and gluconeogenesis, pentose phosphate, TCA cycle and oxidative phosphorylation ^59^, consistent with the biochemical role of AST.

**Fig. 2.**
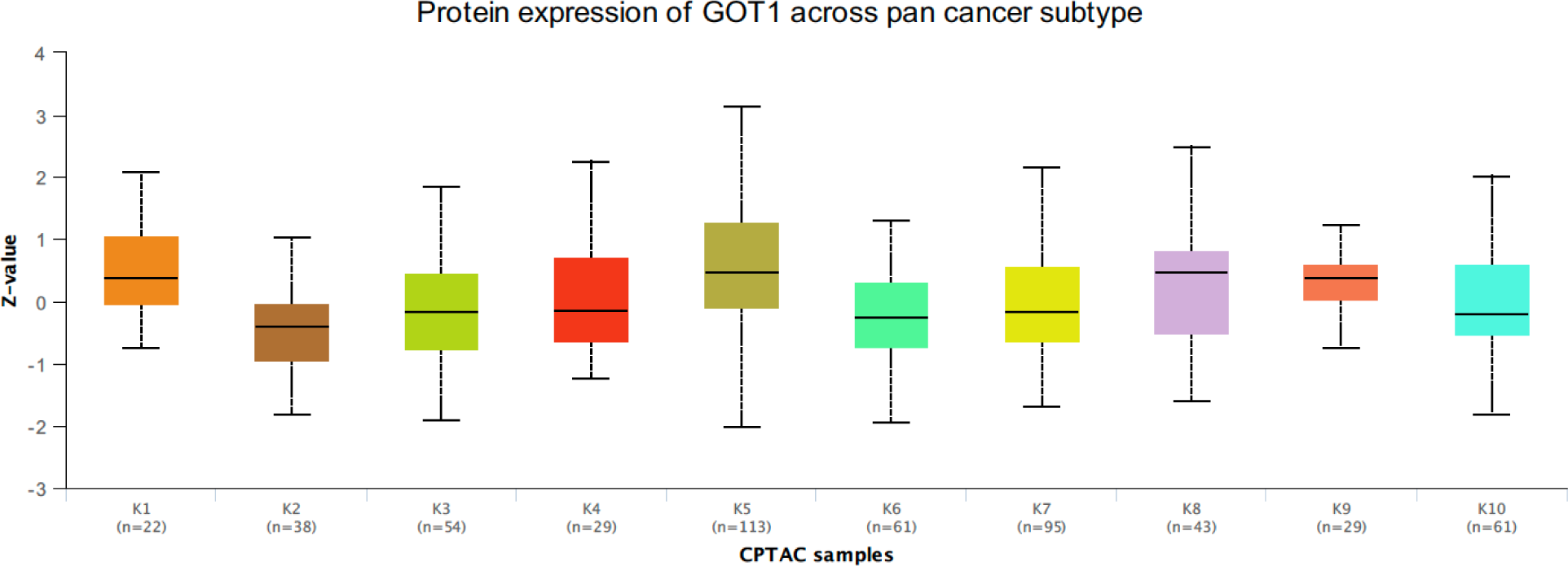
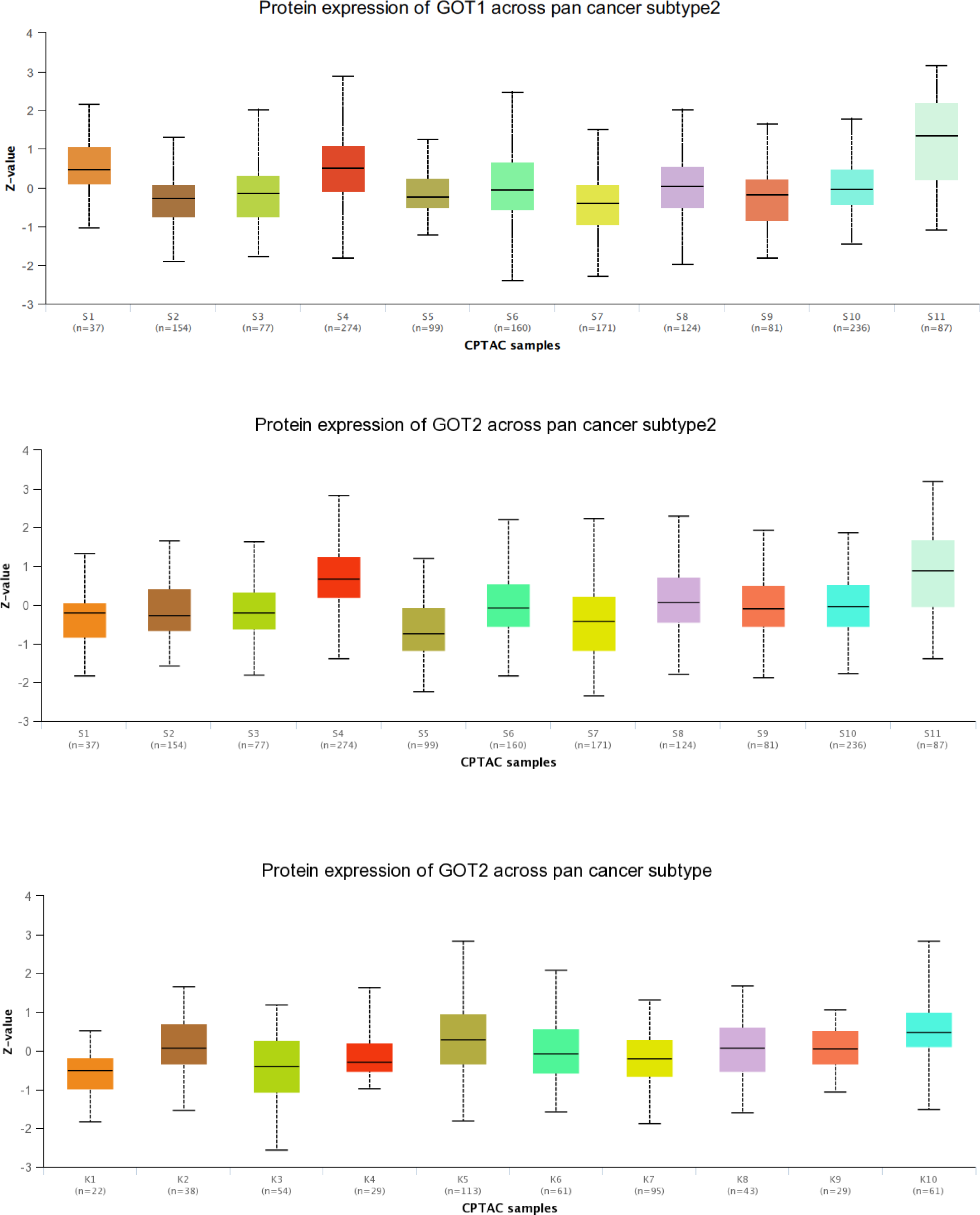
GOT1 and GOT2 expression in previously defined pan-cancer molecular subtypes based on proteogenomic characterization. See Supplementary Tables 1 to 4 for P-values.

In contrast, the cancer subtypes with the least expression of GOT1 were K2 and K6, and S2 and S7. Based on previous studies ^60^, K2 subtypes were associated with MYC targets, K6 with hypoxia, WNT and Epithelial-to-Mesenchymal Transition (EMT). S2 were associated with gene signatures indicating the presence of T-cells and a higher expression of immune checkpoint pathway genes and S7 associated with “axon guidance” and “frizzled binding genes” ^59^.

The top pan-cancer subtypes associated with high expression of GOT2 were S4 and S11 subtypes (also associated with GOT1), and K10 and K5 (also associated with GOT1) subtypes (Fig. 2). The K10 pan-cancer subtype was previously characterized by elevated ER-related proteins and steroid biosynthesis pathway protein, while the K5 subtype was associated with YAP1 and MYC targets, and Ras pathway ^60^.

The pan-cancer subtypes S5 and K3 had the lowest expression of GOT2 transcript levels (Fig. 2). S5 subtype tumors have been associated with B cells, mast cells, eosinophils and elevated expression of complement pathway genes ^59^. In contrast, the K3 subtype was previously associated with hypoxia and EMT signatures. Furthermore, K3 was also enriched with gene signatures for markers of mast cells, macrophages, eosinophils, neutrophils and the complement system ^60^.

### AST transcript levels, tumor prognosis and immune infiltration signatures

We examined the association of GOT1 and GOT2 transcript levels with prognosis in TCGA tumors using patient groups classified as expressing high or low levels of either gene. High expression of GOT1 was significantly associated with poorer survival in LUAD and LAML, and better survival in CESC, KIRC and KIRP (Fig. 3). However, GOT1 expression was not associated with survival in several cancers where these genes were differentially expressed relative to normal tissue.

**Fig. 3.**
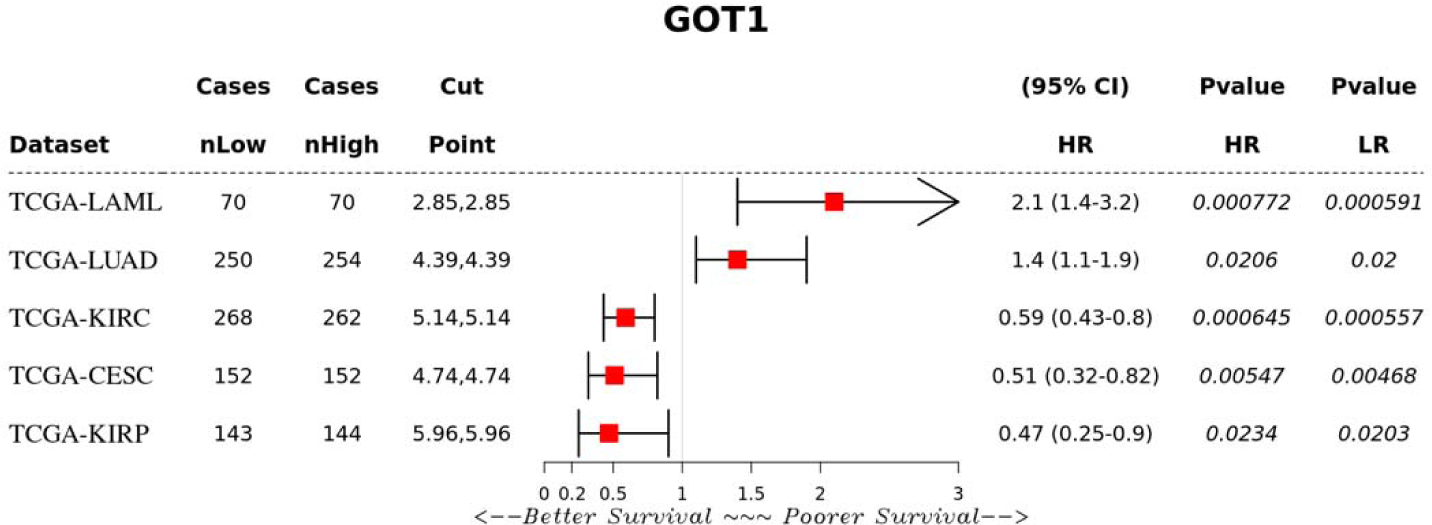
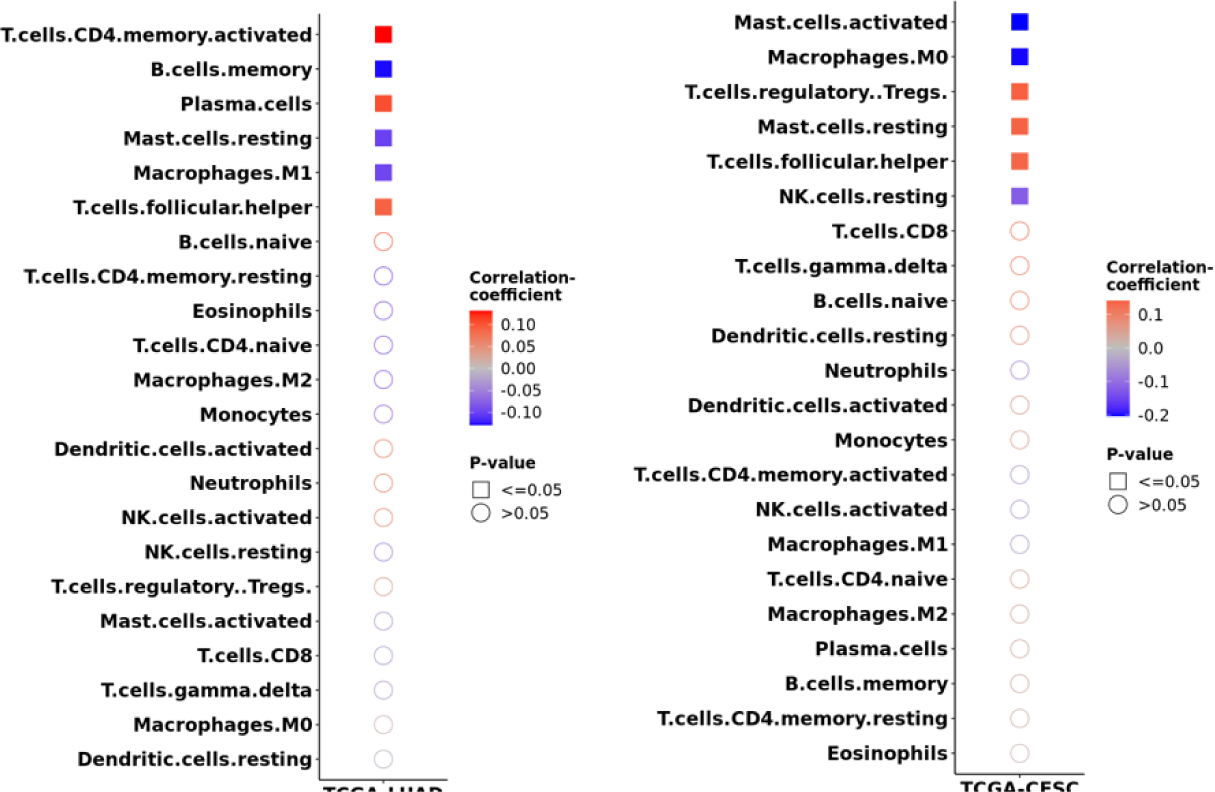
GOT1 expression and associations to overall survival and immune cell infiltration.

To investigate the potential role of tumor infiltrating lymphocytes (TILs) in the distinct association between GOT1 and survival in LUAD and CESC, we examined the correlation between GOT1 expression and TILs based on gene expression signatures. In both tumors, GOT1 expression was positively correlated with follicular helper T-cells. In LUAD, GOT1 was positively correlated with activated CD4+ memory T-cells and plasma cells, and negatively correlated with memory B-cells, resting mast cells and M1 macrophages (Fig. 3). In contrast, GOT1 expression in CESC was positively correlated to regulatory T-cells (Tregs) and resting mast cells, and negatively correlated to activated mast cells, M0 macrophages and Natural Killer (NK) cells (Fig. 3). The positive correlation between GOT1 and Tregs in CESC is consistent with a recent report demonstrating that GOT1 expression promotes the differentiation of Tregs ^74,75^. While Tregs are generally associated with poorer survival outcomes due to their immunosuppressive nature ^76^, high GOT1 expression in CESC was associated with better overall survival (Fig. 3). These results suggest that GOT1 may have context-specific effects on tumor immune responses.

Next, we investigated the association between GOT2 expression and overall survival. High expression level of GOT2 was associated with better overall survival in LIHCs, adrenocortical carcinoma (ACC), KIRP and UCEC, and poorer survival in head and neck cancer (HNSC), mesothelioma (MESO), Brain Lower Grade Glioma (LGG) and Acute Myeloid Leukemia (LAML) (Fig. 4 and Supplementary Fig. 1). GOT2 expression was not associated with survival in several cancers where these genes were differentially expressed relative to normal tissue.

**Fig. 4.**
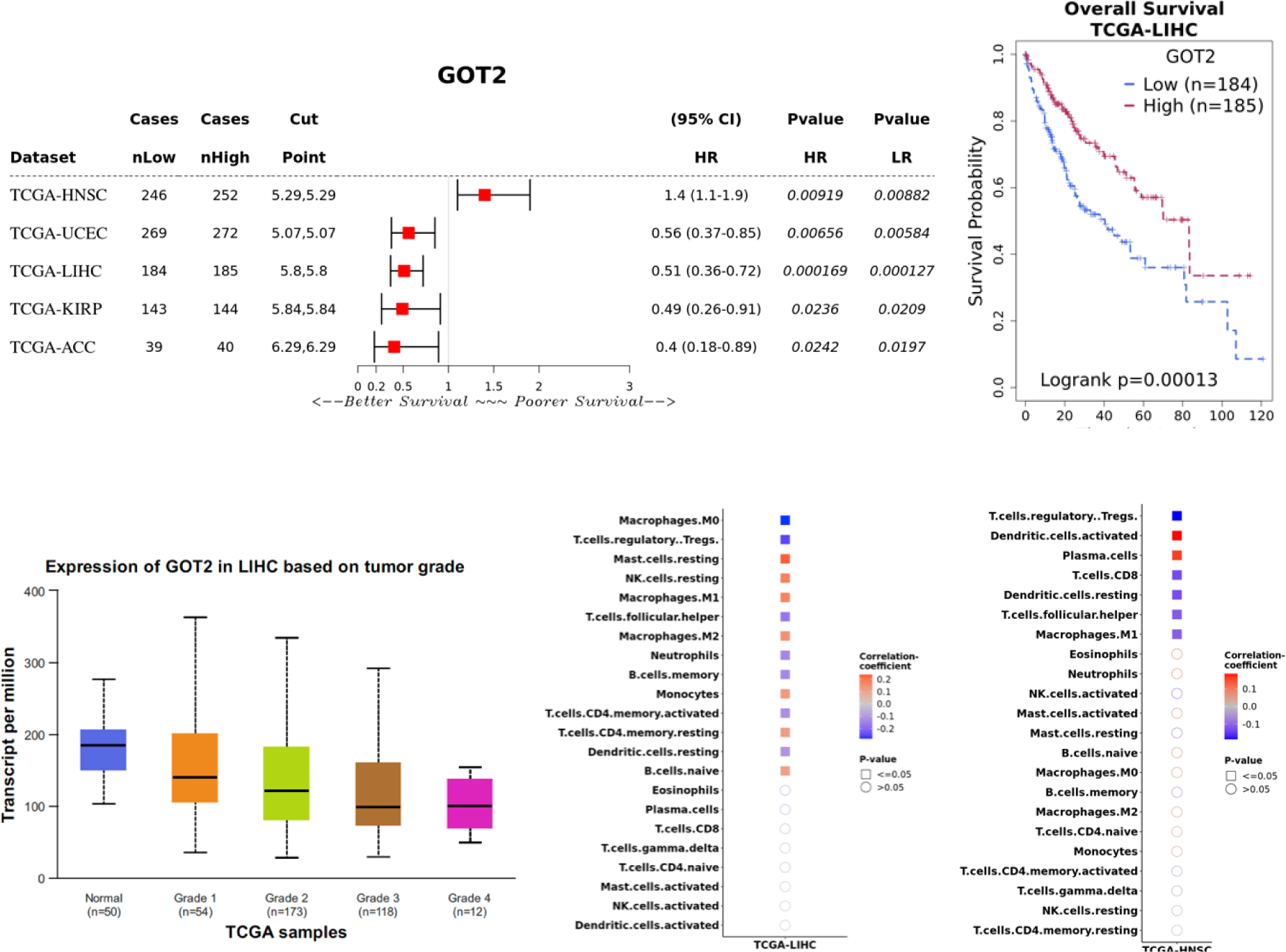
GOT2 expression and associations to overall survival and immune cell infiltration.

The prognostic associations between GOT2 and LIHC are in alignment with genetic studies in which GOT2 knockdown was reported to enhance the proliferation, migration, and invasion of HCC cells in vitro and in mouse models ^73,77^. Furthermore, CAR-T cells engineered to express GOT2 had increased metabolism and anti-tumor activity ^53^. Analysis of inferred TILs in LIHC revealed positive correlation between GOT2 and mast cells, NK cells, M1 and M2 macrophages, monocytes, CD4+ resting T-cells and naïve B-cells. In contrast, GOT2 was negatively correlated with M0 macrophages, Tregs, follicular T-helper, neutrophils, memory B-cells, activated CD4+ T-cells and resting dendritic cells (Fig. 4). Since HNSCs are the only tumors in which a high GOT2 expression was associated with poor overall survival, we assessed the correlation between GOT2 expression and TILs. HNSC tumors had distinct patterns of TILS compared to LIHC and exhibited a positive correlation between GOT2 expression and activated dendritic cells and plasma cells (Fig. 4).

### AST protein-protein networks reveal physical interactions with tumor oncogenes and suppressors

We hypothesized that potential mechanisms underlying GOT1 and GOT2 associations could arise from protein-protein interactions involving other proteins that participate in cancer related molecular pathways. Therefore, we examined public protein-protein interaction data from the BioGrid database for GOT1 and GOT2 interactomes (Supplementary Table 5 and 6). The GOT1 subnetwork from BioGrid contains 269 nodes, of which 263 correspond to distinct proteins and five correspond to small molecules (Supplementary Table 5 and Supplementary Figure 2). In contrast, the GOT2 subnetwork contains 90 proteins and four small molecules (Supplementary Table 6 and Supplementary Figure 3). Interestingly, the top hub in both subnetworks is Parkin, a ubiquitin ligase encoded by the PARK2 gene in which mutations were first found to cause autosomal recessive juvenile parkinsonism ^78,79^. In addition to its role in the survival of neurons^15^, PARK2 is a tumor suppressor ^80^ and has been associated with several types of cancer, including glioblastomas ^81^, lung ^82^, breast^83^, ovarian ^82^, colorectal ^84^ and liver cancers ^85^.

In the GOT1 subnetwork, the presence of EGFR is notable as it is the second most connected gene in this subnetwork (Supplementary Figure 2). EGFR is a well-studied tumor oncogene mutated and/or expressed at high levels in several cancers, especially cancers of the head and neck ^86^, lung ^87^, and the gastrointestinal system ^88–90^. In addition, EGFR is the target of several approved cancer therapies and contributes to cancer drug resistance ^89,91,92^. The GOT1 network also contains GRB2, which binds to EGFR and stimulates KRAS signaling ^93,94^. EGFR and GRB2 are also entry co-factors for the hepatitis B (HBV) ^95^ and C viruses (HCV) ^96^ which contribute to liver cancer ^97,98^. The GOT2 subnetwork includes other critical cancer associated genes: MYC- an important transcription factor overexpressed in many tumors, and PARP1, an enzyme involved in the error-prone DNA repair pathway microhomology-mediated end joining (MMEJ) and a therapeutic target ^99,100^. The protein-protein interactions between both GOT1 and GOT2 with several cancer associated genes provide a potential mechanism through which these genes may be associated with cancer.

To perform an unbiased assessment of potential disease associations between proteins in the GOT1 and GOT2 networks, we analyzed disease enrichments using gene-disease associations from the Disease Gene Network database (DisGeNET) ^66^. GOT1 interaction partners are highly enriched with proteins associated with dermatologic disorders including atopic dermatitis, contact hypersensitivity and acanthosis, and several cancers including adenocarcinoma of the lung and squamous cell carcinoma of the esophagus (Supplementary Table 7). In contrast, GOT2 partners are enriched with cancers of the liver, lung and brain (Supplementary Table 8). We provide detailed list of GOT1 and GOT2 interaction partners that are involved in the enriched diseases in Supplementary Tables 7 and 8.

### AST protein-protein networks are enriched with interaction targets of cancer associated transcription factors and miRNAs

We hypothesized that associations between GOT1 and GOT2 with cancer related biological processes may be mediated through shared protein-protein interactions with transcription factors. Therefore, we assessed whether the protein-protein interactomes of specific transcription factors are enriched with proteins in the GOT1 and GOT2 networks (Supplementary Table 9 and 10). For both GOT1 and GOT2 networks, Estrogen Receptor 1 (ESR1) and Activating Transcription Factor 2 (ATF2)- a driver of tumor aggressiveness ^101^ were ranked in the top 3 transcription factors whose partners were enriched (P << 0.05 in all cases; Supplementary Table 7 and 8). While the transcription factor USF2 (Upstream Stimulatory Factor 2, also associated with several cancers ^102^) was ranked second for GOT1, the overlap between its network and that of GOT2 was ranked much lower (rank = 21; P = 0.002). Notably, the tumor suppressor TP53 was ranked second for the GOT2 network (P = 1.6e-09) but no significant enrichment of its partner proteins was found for GOT1 at P < 0.05. Surprisingly, some of the metabolic enzymes in the GOT2 protein-protein interaction network also interact with P53. Specifically, GAPDH, NQO1, COX17, TX interact with both TP53 and GOT2 (Supplementary Table 8). These interactions could impact cancer progression through several ways. For example, mutant P53 has been shown to prevent the nuclear translocation of GAPDH thereby enhancing glycolysis in cancer cells and inhibiting cell death mechanisms mediated by nuclear GAPDH ^103,104^, NQO1 stabilizes P53 ^105,106^ and TX enhances the binding of P53 to DNA107.

Next, we assessed the GOT1 and GOT2 networks for targets of miRNAs and identified several miRNAs whose targets are enriched in these networks (Supplementary Table 11 and 12). For GOT1, the targets of mir-34a-5p (P = 1.4e-08) and mir-124-3p (P = 4.7e-07) were enriched while for GOT2, mir-4476 (P = 0.00005) and mir-6876-5p (P = 0.00005) targets were enriched. mir-34a-5p is a tumor suppressor and has been associated with several cancers ^108,109^ and diseases such as psoriasis ^110^, MASLD ^111^, MASH ^112^, obesity ^113^, insulin resistance ^114,115^ and osteoarthritis ^116^. This miRNA is a transcriptional target of P53 and regulates P53 directly by targeting TP53 mRNA ^117^ as well as indirectly by targeting the mRNAs encoding negative regulators of P53 including HDM4-a potent negative regulator of P53, resulting in a positive feedback loop ^118^. mir-124-3p is associated with several cancers including HCC, breast and gastric cancers ^119^ as well as neurodegenerative diseases such as Parkinson’s and Alzheimer’s disease ^120^. mir-4476 enriched with GOT2 network members has been associated with gliomas ^121^ and the biological relevance of mir-6876-5p is currently unknown. These results suggest that GOT1 and GOT2 are functionally associated with biological processes in cancer through multiple mechanisms.

## Discussion

In this study, we performed a pan-tissue and pan-cancer analysis of GOT1 and GOT2 transcript and protein levels using publicly available transcriptome and proteomic datasets. GOT1 and GOT2 transcripts were detectable in several tissues though the expression of these genes was particularly high in the skeletal muscle, heart, liver and brain (Fig. 1). Overall, tumors located in tissues that express low levels of GOT1 and GOT2 under normal conditions were characterized by a higher expression of these genes and vice versa. Thus, it appears that relative to their normal corresponding tissue, tumors have a reversed expression of GOT1 and GOT2.

To investigate the potential biological significance of GOT1 and GOT2 expression in tumor biology, we assessed the expression of these genes in previously described pan-cancer molecular subtypes defined using proteomic data ^59,60^. At the regulatory level, our results show that high proteomic levels of GOT1 as well as GOT2 are associated with gene expression signatures of MYC, YAP1 and K-Ras, important players in several cancers. GOT1 and GOT2 tumors are also associated with metabolically similar pan-cancer subtypes characterized by gene expression signatures of the TCA cycle and oxidative phosphorylation. MYC, YAP1 and K-Ras are known to regulate several metabolic processes and have been linked to metabolic reprogramming in many tumors ^122,123^. Thus, the core regulatory programs identified in this study as associated with tumors with high levels of proteomic GOT1 and GOT2 may underlie the enriched metabolic pathway signatures.

Our results suggest that GOT1 and GOT2 expression are associated with distinct immune infiltration patterns that are tissue dependent. Low GOT1 expression was associated with gene expression signatures indicating the presence of T-cells and the expression of immune checkpoint genes, while low GOT2 expression was associated with signatures of various immune cells including mast cells and eosinophils. Among TCGA tumors, high GOT1 expression was associated with poor overall survival in LUAD and LAML and better prognosis in CESC, KIRP and KIRC. In contrast, high GOT2 expression was associated with better prognosis in liver cancer (LIHC), UCEC, KIRP and ACC, and poor prognosis in MESO, HNSC, LGG and LAML. A recent study found that GOT1 promotes the differentiation of Tregs ^74,75^ which provides a possible mechanistic link through which GOT1 may influence prognosis. Although a high GOT1 expression in CESC was associated with increased Treg infiltration (Fig. 3), which generally diminish antitumor immunity, multiple competing factors that interact to influence prognosis could counter this. For example, within CESC tumors, the negative effects of Tregs could be countered by a higher level of T follicular helper cells and resting mast cells, alongside lower levels of activated mast cells and M0 Macrophages, which collectively create a more immunocompetent environment. T-follicular help cells have been associated with favorable outcome in solid organ tumors of non-lymphocytic origin ^124^ including cervical cancer ^125^, activated mast cells were associated with poor outcomes in CESC ^126^ and M0 macrophages have been associated with poor outcomes in several cancers including CESC ^127^.

Our results show that in the case of liver cancer (TCGA LIHC), a high GOT2 expression was associated with an overall immunocompetent tumor environment as several classes of immune cells enriched in high GOT2 expressing tumors have been associated with better prognosis. Specifically, in our results, high GOT2 expression was positively associated with increased infiltration by immune cells that were previously associated with better prognosis including resting mast cells, resting NK cells, M1 macrophages, resting memory CD4 T-cells ^128^ and naïve B-cells ^129^ as well as those that have been associated with poor prognosis in liver cancer such as M2 macrophages ^130^ and monocytes ^131^. A high GOT2 expression was negatively correlated to immune cells associated with poor prognosis including Tregs and M0 macrophages ^132^, as well as to those previously associated with better prognosis – CD4+ memory T-cells ^133^, and dendritic cells ^134^. Thus, even though high GOT2 expression was associated with some lymphocyte subsets that generally have a negative impact on prognosis, the overall prognosis of LIHC patients with high GOT2 expression may have a better outcome. The association between GOT2 with immune infiltrating cells could have implications in the pathology of other diseases accompanied with dysfunctional expression of GOT2 such as MASH and MASLD.

To gain deeper biological insights into potential connections between the biological functions of GOT1 and GOT2, we leveraged experimentally validated and manually curated human protein-protein interactions from the BioGrid database. The direct interaction between GOT1 and GOT2 with known tumor suppressors and oncogenes including EGFR and MYC is notable but further experimental validation of these interactions is needed as many protein-protein interaction data are obtained from noisy, high-throughput methods which presents a limitation of our study. The EGFR-GOT1 interaction in BioGrid is based on the results of a study that performed targeted characterization of EGFR interaction using a mammalian yeast-two-hybrid system, identifying a set of 87 EGFR interacting proteins ^135^ while the MYC-GOT2 interaction was obtained from SILAC immunoprecipitation of the targets of the ubiquitin dependent ATPase Valosin-Containing Protein (VCP) which degrades several proteins including c-Myc, via the ubiquitin-proteasome pathway ^136^. Independent of the protein-protein interaction networks, our analyses found that pan-cancer molecular subtypes expressing high GOT1 and GOT2 were associated with both Ras and MYC gene expression signatures, providing an orthogonal source of evidence for functional associations between GOT1 and GOT2 with Ras (a component of the EGFR signaling cascade) and Myc. The protein-protein interaction data also show that GRB2, an adaptor protein that relays EGFR signals to Ras, interacts with GOT1. Interestingly, the GOT1-GRB2 interaction is among protein-protein interactions that were reported as rewired in colorectal cancer cells expressing the mutant KRAS^G13D^ in which the interaction was only present in cells expressing high levels of mutant KRAS ^137^.

Our analysis of shared protein-protein interactions between transcription factors and GOT1 and 2 revealed key transcription factors including ESR1, TP53, ATF2 and USF2 that have all been associated with various cancers. ESR1 is frequently mutated in estrogen receptor (ER) positive and ER-resistant breast cancers which show increased dependence on glutamine ^138^. Furthermore, GOT1 and GOT2 were found to be critical for the growth of wild-type TP53 expressing cells under glutamine starvation ^139^. ATF2 is a transcription factor that is phosphorylated during starvation of essential amino acids, resulting into increased transcription of amino acid-regulated genes ^140^. Although glutamine is nonessential for normal cells, cancer cells are addicted to glutamine ^141^ and P53 is a critical regulator of adaptation of cancer cells to glutamine deprivation ^142^, a situation created in tumors by the increased utilization of this amino acid.

Thus, the protein-protein interaction networks reinforce the role of GOT1 and GOT2 in glutamine adaptation in cancer.

Finally, we identified the targets of several cancer-associated miRNAs as enriched in genes encoding GOT1 and GOT2 interacting proteins. Among them, we identified mir-34a-5p, a key miRNA in the regulation of P53 ^117,118^ and an emerging cancer therapeutic target ^143^. While a first-in-human Phase I clinical trial of this miRNA was terminated prematurely due to serious adverse immune-mediated events ^144^, the recent development of a fully modified version of mir-34a with enhanced stability, activity and anti-tumor efficacy resulting in complete cures of some mice has ignited increased interest in therapeutic development ^145^.

## Limitations

It is important to note that the work reported in this study is based on analyses of limited public transcriptional and proteomic datasets. The transcription levels of genes are not always directly correlated with the protein levels. The use of bulk cell transcriptome data which does not differentiate expression in cancer cells vs other cells near the tumor as well as the tumor microenvironment is another limitation. The associations between GOT1 and GOT2 with prognosis may be dependent on other properties of the tumors that were not considered in this study. For example the tumor microenvironment, surrounding cell types, mutation or expression status of other genes and patient features including diversity and treatment exposures. In addition to being noisy, experimentally available protein-protein interaction datasets are often collected in a single cell type. It is possible that many experimentally determined interactions are absent in other cell types and maybe highly regulated developmentally, temporally and in different tumors. The analysis of infiltrating immune cells will in future require experimental validation for any associations with the expression of GOT1 and GOT2 as the current study utilized computational techniques.

## Acknowledgements

The Genotype-Tissue Expression (GTEx) Project was supported by the <u>Common Fund</u> of the Office of the Director of the National Institutes of Health, and by NCI, NHGRI, NHLBI, NIDA, NIMH, and NINDS. The data used for the analyses described in this manuscript were obtained from: the <u>GTEx Portal</u> on 05/11/23. The pan-cancer analyses presented in this work were in part based upon data generated by the TCGA Research Network: https://www.cancer.gov/tcga.

## Availability of Data and Materials

The datasets and analytic resources used in this study area available from the public repositories GTEx (https://gtexportal.org) - for gene expression data across multiple tissues, TCGA (https://www.cancer.gov/tcga) - for gene expression data across multiple tumors alongside survival outcomes, CPTAC (https://proteomic.datacommons.cancer.gov/pdc/) – proteomics data from multiple tumors, UCSC Xena (https://xena.ucsc.edu/) - an online tool for exploring multiple cancer datasets, UALCAN (https://ualcan.path.uab.edu/) – a web resource for analyzing multiple public cancer omics datasets including from TCGA and CPTAC, SurvivalGenie (https://bhasinlab.bmi.emory.edu/SurvivalGenie/) – a web platform for survival analysis using gene expression data across multiple tumors in public databases, BioGRID (https://thebiogrid.org/) – a database of protein, genetic and chemical interactions across multiple species, DisGeNET (https://www.disgenet.org/) – a database of gene-disease associations and Enrichr (https://maayanlab.cloud/Enrichr/) – a webserver for performing a wide range of gene set enrichment analyses.

## Supplementary Information

**Supplementary Figure 1:**
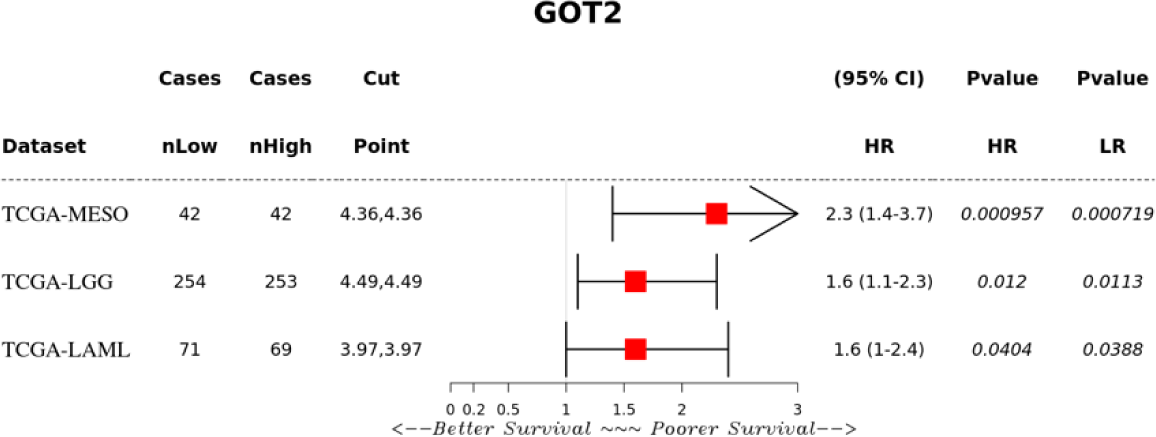

**Supplementary Figure 2:**
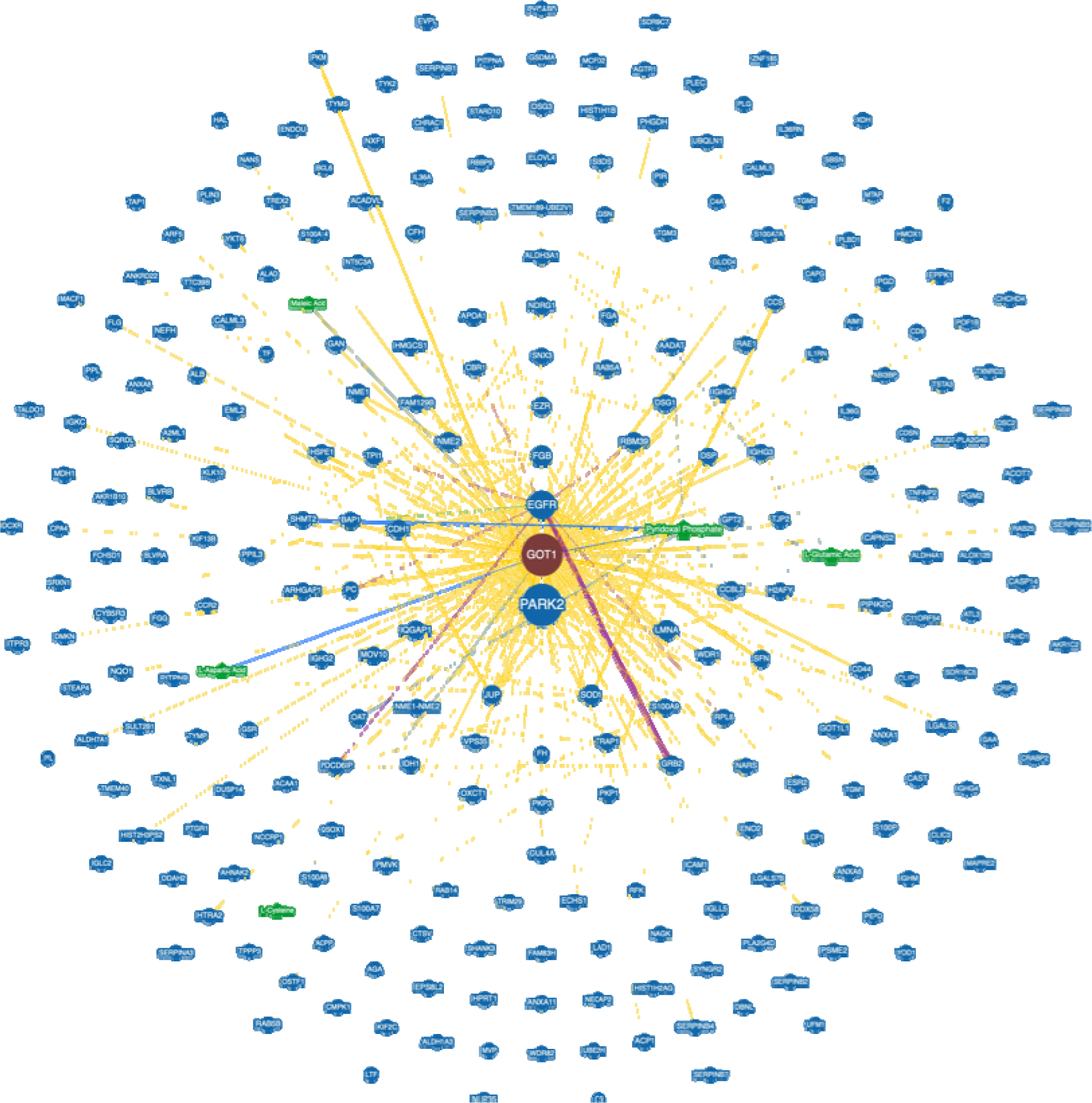
Network diagram of GOT1 interactome from BioGrid

**Supplementary Figure 3:**
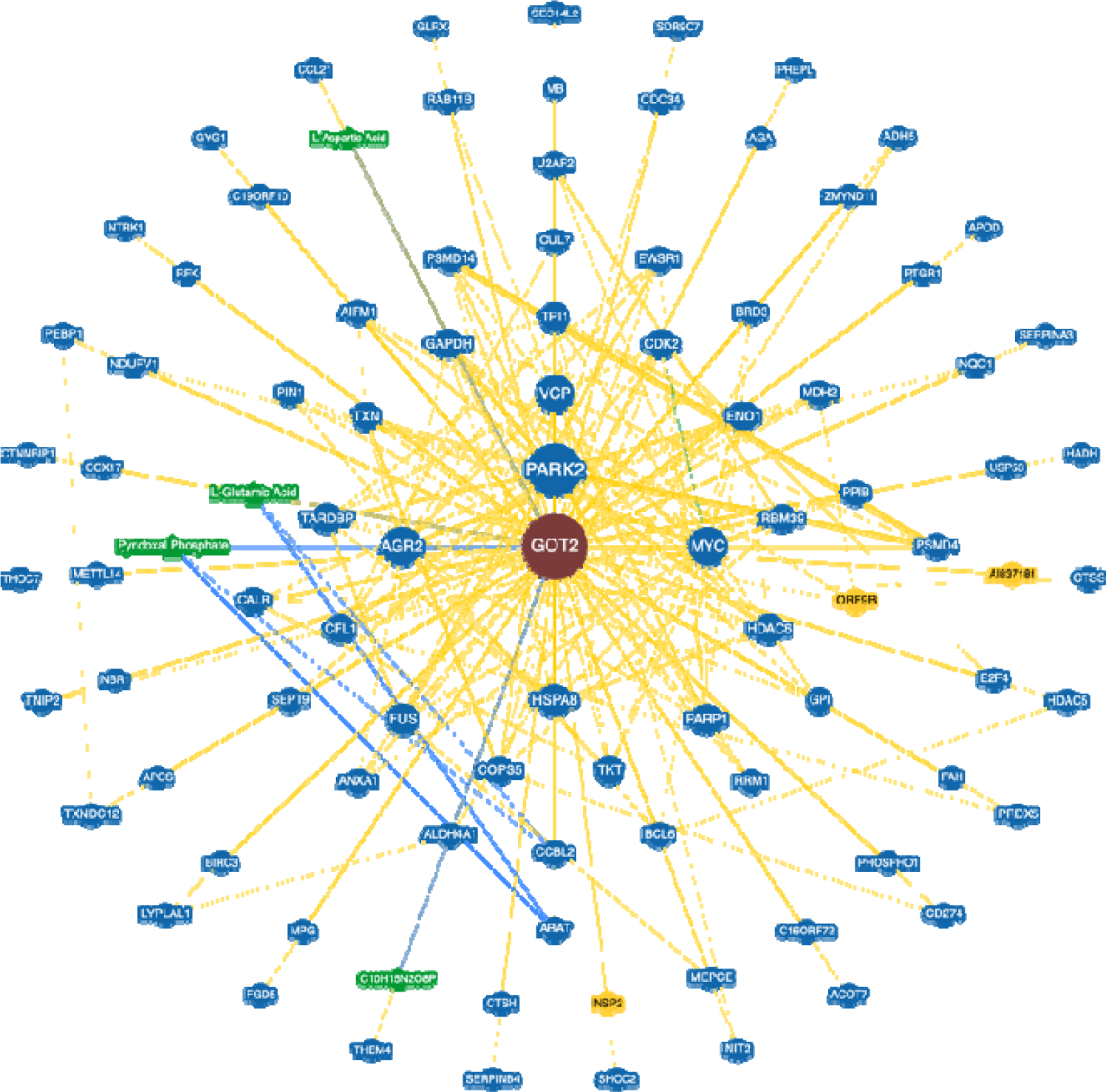
Network diagram of GOT2 interactome from BioGrid.

**Supplementary Table 1:** Protein expression of GOT1 across pan-cancer subtype 2.

| Comparison | Statistical significance |
| --- | --- |
| *S1-vs-S2* | 1.05896551482327E-06 |
| *S1-vs-S3* | 1.242520E-04 |
| *S1-vs-S4* | 5.505592E-01 |
| *S1-vs-S5* | 5.9719789101792E-06 |
| *S1-vs-S6* | 4.646380E-03 |
| *S1-vs-S7* | 3.12188814823019E-08 |
| *S1-vs-S8* | 2.276876E-04 |
| *S1-vs-S9* | 3.96877412611994E-06 |
| *S1-vs-S10* | 4.741356E-04 |
| *S1-vs-S11* | 1.633536E-03 |
| S2-vs-S3 | 2.369850E-01 |
| **S2-vs-S4** | **3.17981524479123E-16** |
| S2-vs-S5 | 5.156640E-01 |
| **S2-vs-S6** | **6.612786E-04** |
| S2-vs-S7 | 7.886367E-02 |
| **S2-vs-S8** | **3.014693E-02** |
| S2-vs-S9 | 9.601194E-01 |
| **S2-vs-S10** | **1.100498E-03** |
| **S2-vs-S11** | **1.29784997001995E-19** |
| **S3-vs-S4** | **1.70534834722379E-06** |
| S3-vs-S5 | 5.170667E-01 |
| S3-vs-S6 | 1.034156E-01 |
| **S3-vs-S7** | **1.648686E-02** |
| S3-vs-S8 | 5.707774E-01 |
| S3-vs-S9 | 3.008542E-01 |
| S3-vs-S10 | 2.604719E-01 |
| **S3-vs-S11** | **6.73739509105303E-14** |
| **S4-vs-S5** | **7.50751698614308E-12** |
| **S4-vs-S6** | **4.085093E-04** |
| **S4-vs-S7** | **1.11359845842929E-19** |
| **S4-vs-S8** | **3.38688743156801E-07** |
| **S4-vs-S9** | **1.09681640140354E-09** |
| **S4-vs-S10** | **1.27986611626931E-07** |
| **S4-vs-S11** | **1.39115723141316E-06** |
| **S5-vs-S6** | **7.523038E-03** |
| **S5-vs-S7** | **2.696215E-02** |
| S5-vs-S8 | 1.450617E-01 |
| S5-vs-S9 | 5.849375E-01 |
| **S5-vs-S10** | **2.045250E-02** |
| **S5-vs-S11** | **7.3084011011793E-18** |
| **S6-vs-S7** | **3.96589539289335E-06** |
| S6-vs-S8 | 2.118035E-01 |
| **S6-vs-S9** | **5.208718E-03** |
| S6-vs-S10 | 4.144923E-01 |
| **S6-vs-S11** | **1.13815282957725E-11** |
| **S7-vs-S8** | **4.173384E-04** |
| S7-vs-S9 | 1.960367E-01 |
| **S7-vs-S10** | **2.59025287743712E-06** |
| **S7-vs-S11** | **3.24352826697589E-22** |
| S8-vs-S9 | 7.912348E-02 |
| S8-vs-S10 | 5.392389E-01 |
| **S8-vs-S11** | **1.24391885617478E-14** |
| **S9-vs-S10** | **1.442506E-02** |
| **S9-vs-S11** | **6.11988917222274E-17** |
| **S10-vs-S11** | **1.37401493343956E-14** |

**Supplementary Table 2:** Protein expression of GOT1 across pan-cancer subtypes.

| *Comparison* | *Statistical significance* |
| --- | --- |
| **K1-vs-K2** | **2.049439E-04** |
| **K1-vs-K3** | **2.904263E-03** |
| **K1-vs-K4** | **4.569760E-02** |
| K1-vs-K5 | 4.507418E-01 |
| **K1-vs-K6** | **2.591916E-04** |
| **K1-vs-K7** | **7.214333E-03** |
| K1-vs-K8 | 1.730756E-01 |
| K1-vs-K9 | 1.705807E-01 |
| **K1-vs-K10** | **3.392581E-02** |
| K2-vs-K3 | 2.357407E-01 |
| K2-vs-K4 | 5.160282E-02 |
| **K2-vs-K5** | **7.64357514696217E-06** |
| K2-vs-K6 | 5.399513E-01 |
| **K2-vs-K7** | **3.553567E-02** |
| **K2-vs-K8** | **1.852669E-03** |
| **K2-vs-K9** | **1.399825E-04** |
| **K2-vs-K10** | **4.617745E-02** |
| K3-vs-K4 | 3.355686E-01 |
| **K3-vs-K5** | **4.622581E-04** |
| K3-vs-K6 | 3.999099E-01 |
| K3-vs-K7 | 4.262346E-01 |
| **K3-vs-K8** | **3.608940E-02** |
| **K3-vs-K9** | **6.304371E-03** |
| K3-vs-K10 | 3.408621E-01 |
| K4-vs-K5 | 6.364198E-02 |
| K4-vs-K6 | 7.893589E-02 |
| K4-vs-K7 | 6.719550E-01 |
| K4-vs-K8 | 3.704872E-01 |
| K4-vs-K9 | 2.313805E-01 |
| K4-vs-K10 | 9.543790E-01 |
| **K5-vs-K6** | **2.05664495503882E-08** |
| **K5-vs-K7** | **4.729602E-04** |
| K5-vs-K8 | 3.155248E-01 |
| K5-vs-K9 | 2.820547E-01 |
| **K5-vs-K10** | **3.846192E-02** |
| **K6-vs-K7** | **3.335331E-02** |
| **K6-vs-K8** | **1.321725E-03** |
| **K6-vs-K9** | **6.47822039763312E-06** |
| K6-vs-K10 | 6.778317E-02 |
| K7-vs-K8 | 9.389489E-02 |
| **K7-vs-K9** | **1.303132E-02** |
| K7-vs-K10 | 7.038927E-01 |
| *K8-vs-K9* | 8.399931E-01 |
| *K8-vs-K10* | 3.145936E-01 |
| *K9-vs-K10* | 1.742075E-01 |

**Supplementary Table 3:** Protein expression of GOT2 across pan-cancer subtype 2.

| *Comparison* | *Statistical significance* |
| --- | --- |
| *S1-vs-S2* | 2.764144E-01 |
| *S1-vs-S3* | 5.428634E-01 |
| ***S1-vs-S4*** | **2.41212357291803E-08** |
| ***S1-vs-S5*** | **1.148701E-02** |
| *S1-vs-S6* | 1.112234E-01 |
| *S1-vs-S7* | 3.247455E-01 |
| ***S1-vs-S8*** | **2.582699E-02** |
| *S1-vs-S9* | 2.769208E-01 |
| *S1-vs-S10* | 1.805810E-01 |
| ***S1-vs-S11*** | **1.11476156919406E-07** |
| *S2-vs-S3* | 5.654427E-01 |
| ***S2-vs-S4*** | **8.49541439613905E-19** |
| ***S2-vs-S5*** | **7.94529466203424E-08** |
| *S2-vs-S6* | 3.946192E-01 |
| ***S2-vs-S7*** | **1.322986E-03** |
| *S2-vs-S8* | 5.268595E-02 |
| *S2-vs-S9* | 8.911707E-01 |
| *S2-vs-S10* | 6.999764E-01 |
| ***S2-vs-S11*** | **1.85594834366474E-09** |
| ***S3-vs-S4*** | **3.14018600300725E-13** |
| ***S3-vs-S5*** | **4.92663722266389E-05** |
| *S3-vs-S6* | 2.074913E-01 |
| ***S3-vs-S7*** | **3.329282E-02** |
| ***S3-vs-S8*** | **3.220853E-02** |
| *S3-vs-S9* | 5.418423E-01 |
| *S3-vs-S10* | 3.613515E-01 |
| ***S3-vs-S11*** | **4.06302656131284E-09** |
| ***S4-vs-S5*** | **9.73843895147848E-32** |
| ***S4-vs-S6*** | **7.28745103435624E-15** |
| ***S4-vs-S7*** | **1.77358363987216E-28** |
| ***S4-vs-S8*** | **4.00507519266657E-12** |
| ***S4-vs-S9*** | **3.32964069691466E-11** |
| ***S4-vs-S10*** | **9.25502650607317E-22** |
| *S4-vs-S11* | 6.060741E-01 |
| ***S5-vs-S6*** | **2.9024869959487E-09** |
| *S5-vs-S7* | **1.928343E-02** |
| *S5-vs-S8* | **7.79326443356387E-12** |
| *S5-vs-S9* | **4.0850753367784E-06** |
| *S5-vs-S10* | **1.83276691644722E-09** |
| *S5-vs-S11* | **1.18975466458995E-18** |
| *S6-vs-S7* | **9.54038773671492E-05** |
| *S6-vs-S8* | 3.000943E-01 |
| *S6-vs-S9* | 5.784134E-01 |
| *S6-vs-S10* | 5.754463E-01 |
| *S6-vs-S11* | **4.88737742534135E-08** |
| *S7-vs-S8* | **8.02086442103514E-07** |
| *S7-vs-S9* | **5.878561E-03** |
| *S7-vs-S10* | **1.294477E-04** |
| *S7-vs-S11* | **1.54654873609547E-14** |
| *S8-vs-S9* | 1.556448E-01 |
| *S8-vs-S10* | 8.031637E-02 |
| *S8-vs-S11* | **1.09129085965805E-06** |
| *S9-vs-S10* | 8.787951E-01 |
| *S9-vs-S11* | **7.23896018928741E-08** |
| *S10-vs-S11* | **2.25109659096499E-09** |

**Supplementary Table 4:** Protein expression of GOT2 across pan-cancer subtypes.

| *Comparison* | *Statistical significance* |
| --- | --- |
| *K1-vs-K2* | **4.882929E-02** |
| *K1-vs-K3* | 6.583243E-01 |
| *K1-vs-K4* | 2.306996E-01 |
| *K1-vs-K5* | **4.082383E-03** |
| *K1-vs-K6* | 2.452687E-01 |
| *K1-vs-K7* | 6.139935E-01 |
| *K1-vs-K8* | 1.967151E-01 |
| *K1-vs-K9* | 2.040442E-01 |
| *K1-vs-K10* | **7.864264E-04** |
| *K2-vs-K3* | **1.074099E-03** |
| *K2-vs-K4* | 4.259403E-01 |
| *K2-vs-K5* | 2.133390E-01 |
| *K2-vs-K6* | 2.094241E-01 |
| *K2-vs-K7* | **1.597597E-02** |
| *K2-vs-K8* | 2.997606E-01 |
| *K2-vs-K9* | 2.307810E-01 |
| *K2-vs-K10* | **3.992451E-02** |
| *K3-vs-K4* | **4.280462E-02** |
| *K3-vs-K5* | **8.08479731226368E-07** |
| *K3-vs-K6* | **2.073307E-02** |
| *K3-vs-K7* | 1.288011E-01 |
| ***K3-vs-K8*** | **1.413581E-02** |
| ***K3-vs-K9*** | **1.102529E-02** |
| ***K3-vs-K10*** | **7.64441023061932E-07** |
| *K4-vs-K5* | 5.893692E-02 |
| *K4-vs-K6* | 8.172209E-01 |
| *K4-vs-K7* | 2.675084E-01 |
| *K4-vs-K8* | 9.517425E-01 |
| *K4-vs-K9* | 8.873739E-01 |
| ***K4-vs-K10*** | **1.124012E-02** |
| ***K5-vs-K6*** | **5.499208E-03** |
| ***K5-vs-K7*** | **1.01760338526256E-05** |
| ***K5-vs-K8*** | **1.303625E-02** |
| ***K5-vs-K9*** | **5.177650E-03** |
| *K5-vs-K10* | 2.579052E-01 |
| *K6-vs-K7* | 2.374458E-01 |
| *K6-vs-K8* | 8.315453E-01 |
| *K6-vs-K9* | 8.981846E-01 |
| ***K6-vs-K10*** | **1.040275E-03** |
| *K7-vs-K8* | 1.671853E-01 |
| *K7-vs-K9* | 1.562240E-01 |
| ***K7-vs-K10*** | **1.31289610484227E-05** |
| *K8-vs-K9* | 9.204135E-01 |
| ***K8-vs-K10*** | **2.240220E-03** |
| ***K9-vs-K10*** | **1.031038E-03** |

**Supplementary Table 5:** GOT1 protein-protein interactions from BioGrid

**Supplementary Table 6:** GOT2 protein-protein interactions from BioGrid

**Supplementary Table 7:** Top 10 significant p-values and q-values for GOT1 PPis based on DisGeNET.

| Table of top 10 significant p-values and q-values for GOT1 PPis based on DisGeNET |  |  |  |
| --- | --- | --- | --- |
| term | p-value | q-value | overlap_genes |
| Dermatologic disorders | 5.26 E-16 | 2.38 E-12 | [FLG, IL1RN, FH, CALML5, ALOX12B, PLG, DMKN, EGFR, TGM1, S100A7A, NME1-NME2, POF1B, LMNA, HMOX1, IL36RN, NANS, TGM5, TGM3, DSP, CDSN, DDX58, NME2, NME1, DSG1, DSG3, CD44, IVL, DSC2, S100A8, BLVRA, S100A7, PLEC] |
| Neoplasm Metastasis | 8.76 E-14 | 1.99 E-10 | [IL1RN, STEAP4, SDR9C7, ENO2, ICAM1, PYCARD, LGALS3, CDH1, TRIM29, TAP1, ALDH3A1, MTAP, CLIP1, AGTR1, CMPK1, PKP1, HPRT1, PKP3, EZR, S100A9, DSC2, S100A8, S100A7, MACF1, CFH, IQGAP1, NDRG1, NME1- |
|  |  |  | NME2, CYB5R3, HMOX1, SFN, S100A14, TRAP1, CDSN, IDH1, ARHGAP24, ESR2, TF, HAL, AKR1B10, BCL6, ALB, GRB2, RAB5A, BAP1, SERPINA3, CLIC3, ARHGAP1, TNFAIP2, S100A7A, ANXA6, QSOX1, PHGDH, TMEM189-UBE2V1, ACP1, CCR2, DSP, SERPINB4, CBR1, ANXA1, TPI1, SERPINB2, DBNL, ELOVL4, DDX58, NME2, APOA1, TYK2, F2, SERPINB5, NME1, ALDH1A3, PKM, KIF2C, LCP1, DSG3, ACPP, CD44, PLEC, LTF, FH, MVP, CAPG, PLG, TYMS, SBSN, CRIP1, EGFR, TYMP, NXF1, RAB25, ABI3BP, LMNA, XDH, NQO1, LGALS7B, GSN, TXNRD2, AKR1C2, HSPE1, ...] |
| Squamous cell carcinoma | 9.35 E-13 | 1.41 E-09 | [IL1RN, DMKN, ICAM1, PYCARD, TGM1, S100A7A, LGALS3, CDH1, TRIM29, ANXA8, NANS, TGM3, CCR2, SERPINB3, DSP, SERPINB4, ANXA1, TPI1, SERPINB2, NME2, TALDO1, SERPINB5, NME1, MTAP, PKM, PKP1, DSG1, DSG3, EZR, MAPRE2, S100A9, CD44, IVL, DSC2, S100A8, S100A7, FLG, CRABP2, ALOX12B, PLG, TYMS, PPL, NDRG1, EGFR, TYMP, SNX3, NME1-NME2, HMOX1, SFN, S100A14, NQO1, LGALS7B, GSN, JUP, IDH1, TREX2, GSR, AKR1C2, KLK10, ARHGAP24, ESR2, SOD1, HAL, AKR1B10] |
| Adenocarcinoma of lung (disorder) | 3.52 E-12 | 4.00 E-09 | [CRABP2, CFH, ARHGAP1, CAPG, ENO2, TYMS, NDRG1, EGFR, ICAM1, S100A7A, CDH1, LMNA, UBQLN1, HMOX1, SFN, S100A14, TGM5, XDH, SERPINB3, SERPINB4, NQO1, CBR1, ANXA1, TPI1, GSN, IDH1, APOA1, ARHGAP24, ESR2, NME1, SOD1, ALDH3A1, MTAP, TF, HAL, AKR1B10, PKM, ALB, CD9, BLVRB, GRB2, HPRT1, PKP3, EZR, S100A9, RAB5A, BAP1, CD44] |
| Contact hypersensitivity | 1.02 E-11 | 8.87 E-09 | [SERPINB3, FLG, SERPINB4, NQO1, GSR, AKR1C2, SOD1, AKR1B10, PIR, HMOX1, KIF2C, CD44, S100A8] |
| Psoriasis | 1.59 E-11 | 8.87 E-09 | [FLG, IL1RN, CALML5, PLG, CTSV, EGFR, ICAM1, C3, TGM1, S100A7A, NME1-NME2, C4A, NXF1, ANXA6, HMOX1, IL36RN, S100A14, TGM5, TGM3, CCR2, SERPINB3, CAST, SERPINB4, SERPINB1, CDSN, DDX58, NME2, IL36G, TAP1, APOA1, TYK2, SERPINB8, NME1, DSG1, S100A9, IVL, S100A8, S100A7] |
| Squamous cell carcinoma of esophagus | 1.63 E-11 | 8.87 E-09 | [IL1RN, CRABP2, TNFAIP2, SDR9C7, PLG, IQGAP1, TYMS, SBSN, PPL, EVPL, NDRG1, EGFR, TYMP, RAB25, CDH1, HMOX1, SFN, S100A14, NEFH, TGM3, SERPINB3, DSP, TRAP1, NQO1, ANXA1, TPI1, SERPINB2, AKR1C2, SERPINB5, ARHGAP24, ESR2, NME1, MTAP, PKM, PSME2, BLVRB, GRB2, EZR, ALDH7A1, BAP1, CD44, DSC2, S100A8] |
| Adenocarcinoma | 1.92 E-11 | 8.87 E-09 | [IL1RN, ICAM1, PYCARD, LGALS3, CDH1, PHGDH, ANXA8, ACP1, SERPINB3, DSP, SERPINB4, ANXA1, TPI1, |
|  |  |  | SERPINB2, NME2, APOA1, SERPINB5, NME1, ALDH1A3, MTAP, PKM, AGTR1, PSME2, DSG3, ACPP, S100A9, CD44, DSC2, S100A8, PLEC, CRABP2, CFH, CAPG, PLG, IQGAP1, TYMS, NDRG1, EGFR, TYMP, NME1-NME2, IGKC, RAB25, LMNA, HMOX1, SFN, NQO1, GSN, IDH1, GSR, AKR1C2, KLK10, ARHGAP24, ESR2, SOD1, AKR1B10, ALB, CD9, S100P] |
| Acanthosis | 1.93<br>E-11 | 8.87<br>E-09 | [TGM1, CAST, IL1RN, JUP, DSG1, ALOX12B, IL36RN, SERPINB7, SERPINB8, EGFR] |
| Primary malignant neoplasm of lung | 1.96<br>E-11 | 8.87<br>E-09 | [IL1RN, ENO2, ICAM1, PYCARD, TGM1, S100A7A, LGALS3, CDH1, ACP1, CCR2, SERPINB3, DSP, SERPINB4, CBR1, ANXA1, SERPINB2, NME2, TAP1, PTGR1, SERPINB5, NME1, ALDH1A3, MTAP, PKM, AGTR1, KIF2C, HPRT1, PKP3, DSG3, EZR, ALDH7A1, CD44, S100A8, LTF, TSTA3, CRABP2, CFH, SHMT2, MVP, ALOX12B, PLG, TYMS, SBSN, NDRG1, EGFR, TYMP, NXF1, RAB25, ABI3BP, LMNA, HMOX1, SFN, S100A14, XDH, NQO1, RBM39, GSN, ZNF185, TXNRD2, ARHGAP24, ESR2, SOD1, CUL4A, AKR1B10, ALB, CD9, GRB2, RAB5A, BAP1] |

**Supplementary Table 8:**
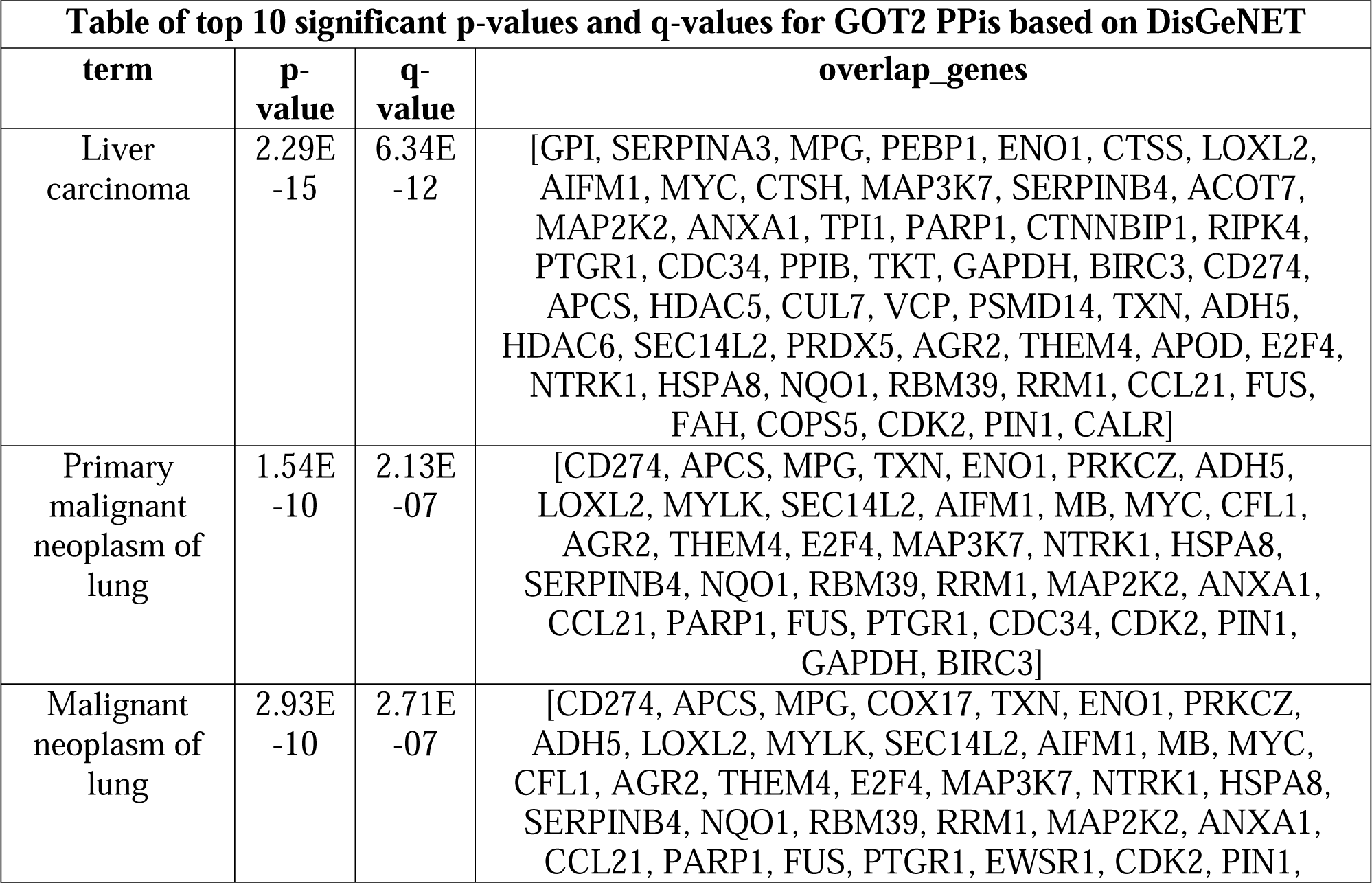
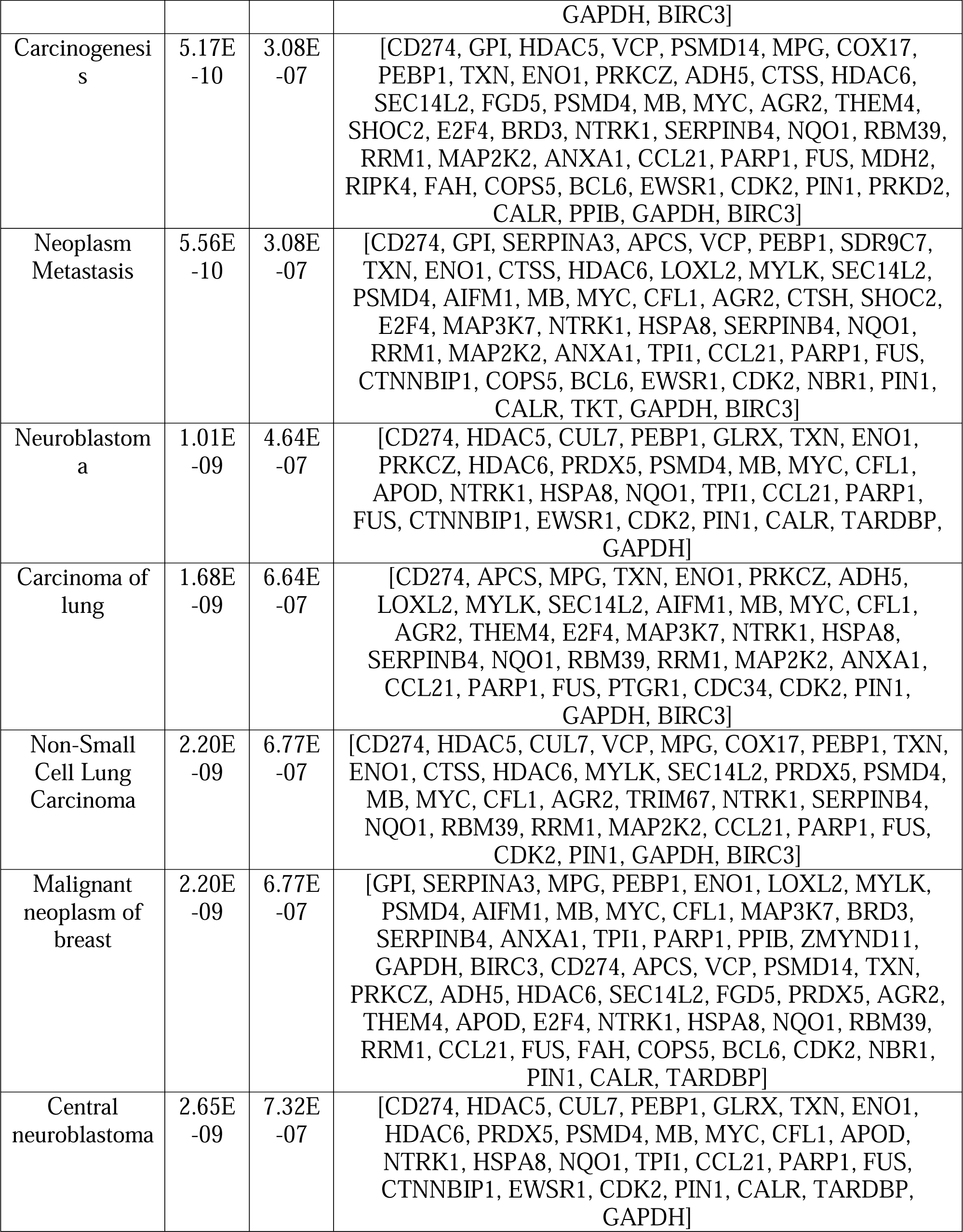
Top 10 significant p-values and q-values for GOT2 PPis based on DisGeNET.

**Supplementary Table 9:**
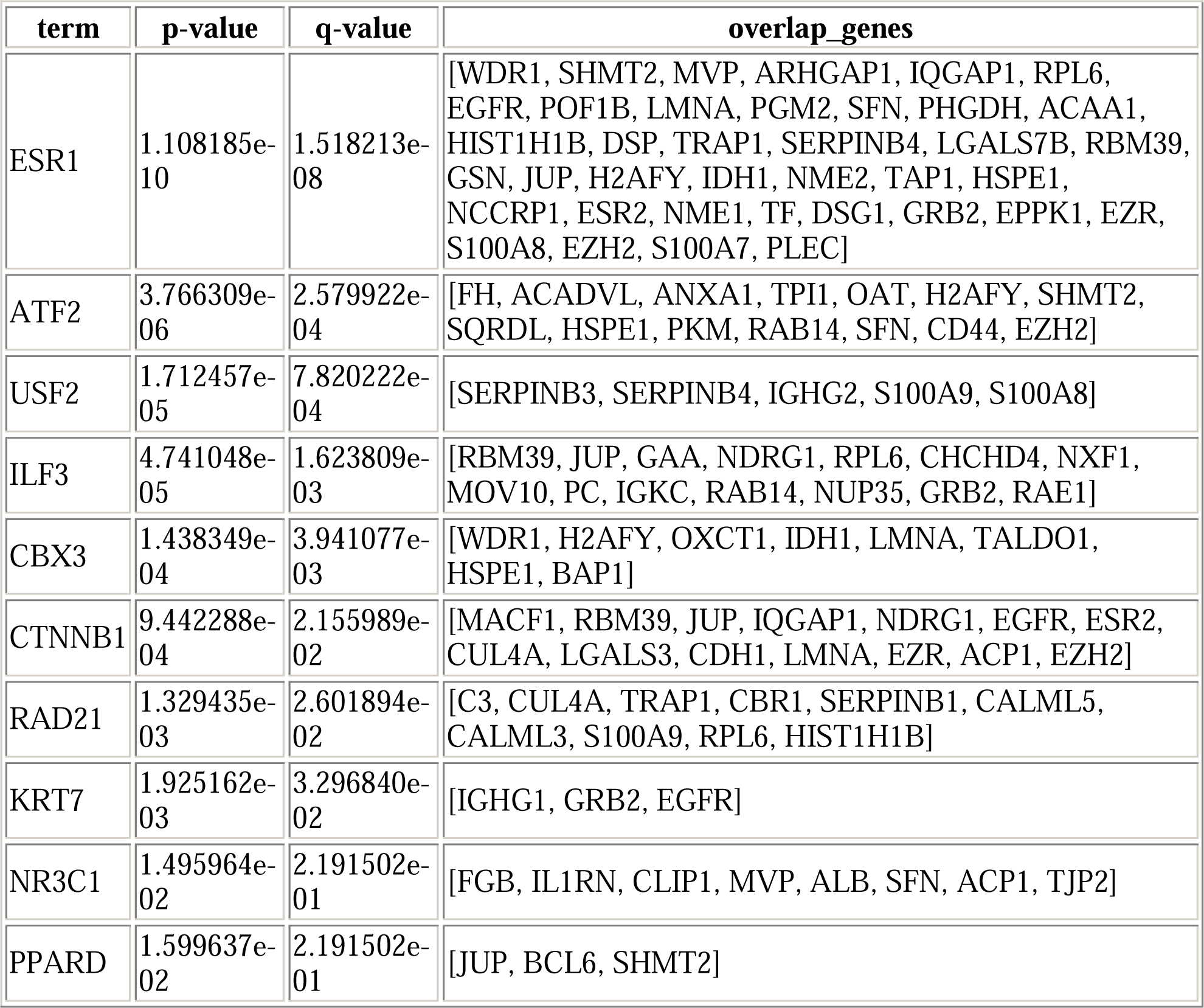
Top 10 significant p-values and q-values for GOT1 Transcription Factor PPIs.

**Supplementary Table 10:** Top 10 significant p-values and q-values for GOT2 Transcription Factor PPIs.

| term | p-value | q-value | overlap_genes |
| --- | --- | --- | --- |
| ESR1 | 5.684151e-13 | 1.051568e-10 | [HDAC5, VCP, MPG, ABAT, ENO1, PRKCZ, AIFM1, U2AF2, MYC, CFL1, SHOC2, APOD, HADH, HSPA8, SERPINB4, RBM39, PARP1, FUS, MDH2, COPS5, EWSR1, CDK2, TKT, GAPDH, EZH2] |
| TP53 | 1.625568e-09 | 1.503650e-07 | [NTRK1, HSPA8, NQO1, HDAC5, CUL7, VCP, PARP1, COX17, TXN, ADH5, COPS5, PSMD4, EWSR1, MYC, CDK2, PIN1, ZMYND11, GAPDH] |
| ATF2 | 2.586069e-08 | 1.594743e-06 | [PRDX5, ANXA1, TPI1, MDH2, CFL1, CDK2, ENO1, CALR, PPIB, GAPDH, EZH2] |
| HIF1A | 3.067008e-06 | 1.218770e-04 | [HDAC5, VCP, COPS5, CDC34, PARP1, MYC, HDAC6] |
| EP300 | 3.822734e-06 | 1.218770e-04 | [BCL6, PARP1, MPG, EWSR1, TNIP2, MYC, CDK2, PIN1, ENO1, GAPDH, PARK2, HDAC6] |
| CTNNB1 | 4.434997e-06 | 1.218770e-04 | [HSPA8, RBM39, COPS5, PARP1, FUS, MYC, CTNNBIP1, CDK2, PIN1, HDAC6, EZH2] |
| HCFC1 | 4.611563e-06 | 1.218770e-04 | [HDAC5, HSPA8, USP53, CDK2, E2F4, MAP3K7] |
| TRIM28 | 7.901592e-06 | 1.827243e-04 | [HDAC5, COPS5, CDC34, PARP1, MYC, MEPCE, CDK2, E2F4] |
| TARDBP | 1.087200e-05 | 2.234799e-04 | [VCP, U2AF2, FUS, PARK2, HDAC6] |
| POLR2A | 2.484885e-05 | 4.597037e-04 | [PARP1, EWSR1, FUS, U2AF2, MYC, CDK2, PIN1, EZH2] |

**Supplementary Table 11:** Top 10 significant p-values and q-values for GOT1 enriched miRNAs miRTarBase 2017.

| Term | p-value | q-value | Overlap_genes |
| --- | --- | --- | --- |
| hsa-miR-34a-5p | 1.36E-08 | 0.000028 | [FH, MCFD2, CRABP2, ARHGAP1, CAPG, TYMS, NDRG1, SNX3, CASP14, SYNGR2, SFN, HIST1H2AG, EPS8L2, RAE1, HIST1H1B, DSP, SERPINB1, YOD1, MOV10, MTAP, DDAH2, AGTR1, CMPK1, PLIN3, GRB2, KIF2C, S100P, HPRT1, SHANK3, CD44, TJP2] |
| hsa-miR-124-3p | 4.70E-07 | 0.000487 | [FLG, ACADVL, CLIC3, GDA, SHMT2, MVP, ARHGAP1, IQGAP1, FAM129B, PYCARD, TGM1, FAM83H, CHRAC1, SYNGR2, TRIM29, LMNA, ANXA6, HMOX1, RBBP9, PGM2, QSOX1, ANXA8, YKT6, CAST, SERPINB2, DBNL, GSN, JUP, TREX2, AHNAK2, ANXA11, TYK2, MOV10, NECAP2, BCL6, GAN, AGTR1, PLIN3, PKP1, GRB2, KIF2C, TJP2, PLEC] |
| hsa-miR-1-3p | 9.78E-05 | 0.067616 | [ACADVL, ECHS1, OAT, ATL3, SRXN1, CAPG, EGFR, CHRAC1, OXCT1, HMOX1, PGM2, SFN, OSTF1, HIST1H1B, CAST, RBM39, ZNF185, JUP, AHNAK2, F2, PGD, SERPINB5, MOV10, PIR, KIF2C, LCP1, CD44] |
| hsa-miR-3659 | 2.19E-04 | 0.113857 | [PKM, TXNL1, VPS35, PIP4K2C, SHANK3, PLEC] |
| hsa-miR-4792 | 3.95E-04 | 0.15377 | [NME1-NME2, CRABP2, GAN, GSR, NME2, CPA4, PLEC] |
| hsa-miR-1304-5p | 4.45E-04 | 0.15377 | [PYCARD, CHCHD4, FAM83H, ALDH1A3, GSN, HMGCS1, FGG, A2ML1] |
| hsa-miR-4443 | 5.92E-04 | 0.175463 | [RAB5B, DCXR, HMOX1, SDR9C7, TRIM67, PLEC, FAM129B] |
| hsa-miR-30a-5p | 8.04E-04 | 0.184444 | [CAST, UBE2H, ANXA1, TPI1, FAHD1, JUP, IDH1, TNFAIP2, SERPINB5, EGFR, ESR2, SNX3, NT5C3A, UFM1, PKM, RAB14, WDR82, CDH1, PLIN3, LCP1, CD44] |
| hsa-miR-4430 | 8.89E-04 | 0.184444 | [RAB5B, TMEM40, F2, SNX3, PYCARD, NARS, MTAP, CCS, PKM, NAGK, GAN, HMOX1, PLIN3, QSOX1] |
| hsa-miR-3652 | 8.89E-04 | 0.184444 | [RAB5B, TMEM40, F2, SNX3, PYCARD, NARS, MTAP, CCS, PKM, NAGK, GAN, HMOX1, PLIN3, QSOX1] |

**Supplementary Table 12:** Top 10 significant p-values and q-values for GOT2 enriched miRNAs miRTarBase 2017.

| Term | p-value | q-value | Overlap_genes |
| --- | --- | --- | --- |
| **hsa-miR-4476** | **0.000051** | **0.03613** | **[ALDH4A1, HDAC5, TPI1, CFL1, SDR9C7, CALR, TRIM67]** |
| **hsa-miR-6876-5p** | **0.000051** | **0.03613** | **[ALDH4A1, HDAC5, TPI1, CFL1, SDR9C7, CALR, TRIM67]** |
| hsa-miR-1301-3p | 0.000408 | 0.181755 | [VCP, PARP1, MDH2, CFL1, TXN, MYLK] |
| hsa-miR-4254 | 0.000511 | 0.181755 | [METTL14, CDC34, SDR9C7, PRKD2] |
| mmu-miR-127-3p | 0.001021 | 0.241095 | [GPI, BCL6] |
| mmu-miR-449a-5p | 0.001186 | 0.241095 | [MYC, PIN1, RAB11B] |
| hsa-miR-615-3p | 0.001244 | 0.241095 | [HSPA8, TPI1, CDC34, PARP1, EWSR1, FUS, MDH2, U2AF2, NBR1, E2F4, ENO1, GAPDH] |
| mmu-miR-34c-5p | 0.001633 | 0.241095 | [BCL6, PIN1, RAB11B] |
| hsa-miR-873-5p | 0.001847 | 0.241095 | [PRDX5, TPI1, CFL1, PIN1, ADH5, RAB11B] |
| hsa-miR-6859-5p | 0.001937 | 0.241095 | [VCP, CFL1, CDK2, PIN1, TRIM67] |

